# Cell Adhesion-Dependent Biphasic Axon Outgrowth Elucidated by Femtosecond Laser Impulse

**DOI:** 10.1101/2021.10.10.463848

**Authors:** Sohei Yamada, Kentarou Baba, Naoyuki Inagaki, Yoichiroh Hosokawa

**Author notes:** These authors contributed equally (Sohei Yamada, Kentarou Baba). Corresponding authors: Sohei Yamada, and Yoichiroh Hosokawa.

## Abstract

Axon outgrowth is promoted by the mechanical coupling between F-actin and adhesive substrates via clutch and adhesion molecules in an axonal growth cone. In this study, we utilized a femtosecond laser-induced impulse to break the coupling between the growth cone and the substrate, enabling us to evaluate the strength of the binding between the growth cone and a laminin on the substrate, and also determine the contribution of adhesion strength to neurite outgrowth and traction force for the outgrowth. We found that the adhesion strength of axonal L1 cell adhesion molecule (L1CAM)-laminin binding increased with the laminin density on the substrate. In addition, fluorescent speckle microscopy revealed that the retrograde flow of F-actin in the growth cone was dependent on the laminin density such that the flow speed reduced with increasing L1CAM-laminin binding. However, neurite outgrowth and the traction force did not increase monotonically with increased L1CAM-laminin binding but rather exhibited biphasic behavior, in which the outgrowth was suppressed by excessive L1CAM-laminin binding. Our quantitative evaluations suggest that the biphasic outgrowth is regulated by the balance between traction force and adhesion strength. These results imply that adhesion modulation is key to the regulation of neurite outgrowth.

## Introduction

During neuronal development, axons elongate and form functional connections with other neurons and relevant cells. The axonal migration is an essential process to regulate formation and regeneration of neuronal network. The previous studies proposed that extracellular guidance cue (chemical ligands) induces the axon outgrowth^1,2^. The growth cone located at the tip of an elongating axon senses chemical ligands in the external environment and undergoes directional migration^1–3^. In addition, growth cone senses a rich source of the extracellular matrix (ECM) to facilitate directional migration. An adhesion coupling between the growth cone and its substrate induces rapid migration and axon outgrowth.

The traction force underlying growth cone migration is regulated by modulation of the coupling efficiency between actin filament (F-actin) retrograde flow and adhesive substrates via clutch and cell adhesion molecules^4,5^. Thus, the traction force transmitted to the substrate through the F-actin-adhesion coupling promotes axon outgrowth^1,6^.

We previously identified shootin1a and cortactin as clutch molecules for growth cone migration^7,8^. These molecules mediate the linkage between F-actin retrograde flow and the cytoplasmic domain of L1 cell adhesion molecule (L1CAM)^9^. The extracellular domain of L1CAM interacts with adhesive ligands such as laminins in the extracellular matrix^10–12^. L1CAM linked to the F-actin flow undergoes gripping (stop) and slipping (retrograde flow) on the substrate. It is the balance between these grip and slip states that regulates growth cone migration^12^.

Our investigation focuses on the mechanism by which the interaction between L1CAM and laminin generates force. The challenge is to quantify key processes of growth cone migration, for which the adhesion strength via L1CAM-laminin binding is fundamental. However, it is difficult to quantify adhesion strength by conventional methods. For instance, the shear flow assay^13,14^ is not suitable for evaluating local adhesion at the interface between a growth cone and the substrate, whereas it is difficult to apply single-cell force spectroscopy^15,16^ to evaluate force without disturbing the adhesion required for axon outgrowth. The adhesion strength of migrating growth cone was previously evaluated form a detachment by micromanipulation with calibrated glass needles^17^. However, these methods are not suitable for the evaluation of adhesion strength at the growth cone level. Therefore, the relationship between adhesion strength and axon outgrowth has not been comprehensively elucidated. We thus developed a method utilizing a femtosecond laser-induced impulsive force, which we used to quantify adhesions between leukocytes and endothelial cells, among epithelial cells, and between neurons and mast cells^18,19^.

A near-infrared femtosecond laser focused through an objective lens into a water solution generates stress and shock waves at the laser focal point. Vogel and his co-workers have observed these generation processes and the propagation dynamics, and confirmed that the femtosecond laser pulse generates stress waves with the smallest photon energy and confines the propagation in the smallest area compared to other pulsed lasers^20,21^. Referring the experimental results, we construct a simplified model to explain the action of these waves to small particles near the laser focal point and confirmed the reliability with experiments. Based on these results, the action is approximated that these waves propagate out spherically and act as an impulsive force on nearby cells. As the force is localized to a diameter of 1–10 µm and breaks intercellular adhesions at a single-cell resolution, we considered the impulse could be used to assess adhesion of the growth cone.

We also developed an original method to quantify the magnitude of the impulsive force by using atomic force microscopy (AFM)^22^, enabling us to quantify the strength of intercellular adhesions on the basis of the force needed to break the connection^18,19,23^. In this method, the force is estimated from vibration of the AFM cantilever which placed instead of the cell samples. The vibration was analyzed with taking propagations of shock and stress waves into account, and then the total force generated at the laser focal point was estimated as a function of the laser pulse energy. We can calibrate the force to break adhesion of the growth cone by applying this estimation to the geometrical relation between the laser focal point and the growth cone to break the adhesion.

In this work, we applied our previously established methods for generating laser-induced impulsive force^7,12^ to investigate the contribution of L1CAM-laminin binding to axon outgrowth. The specific interaction between laminin and L1CAM was confirmed by L1CAM knockdown in neurons. The strength of L1CAM-mediated adhesion was confirmed to be dependent on the density of laminin on the substrate. In addition, we used fluorescent speckle microscopy to observe the motions of F-actin and L1CAM in the axonal growth cone and then further assessed the contribution of L1CAM-laminin binding to F-actin-substrate coupling. These results were compared with axon outgrowth and traction force for the outgrowth as a dependence of laminin density on the substrate. We propose a mechanism of neurite outgrowth that depends on the density of laminin on the substrate, identifying L1CAM–laminin binding as a key regulator of this process.

## Results

### Adhesion breaking by a femtosecond laser-induced impulsive force

Hippocampal neurons cultured for 3 days on a glass-bottom dish coated with 10 µg/ml laminin were placed on an inverted microscope equipped with a femtosecond laser irradiation system. The single-shot femtosecond laser pulses were focused in the vicinity of axonal growth cones to assess the adhesion breaking threshold (Fig. 1A). The force was estimated by measuring the distance from the growth cone at which the laser pulse broke the adhesion to the substrate. For example, the laser with a pulse energy of 700 nJ was initially focused at a position 20 µm from a targeted growth cone. After the first pulse irradiation, the focal point was moved closer to the target in 5 µm steps via an electrical microscope stage until adhesion was broken; the distance between the growth cone and the final laser focal position was recorded.

**Figure 1.**
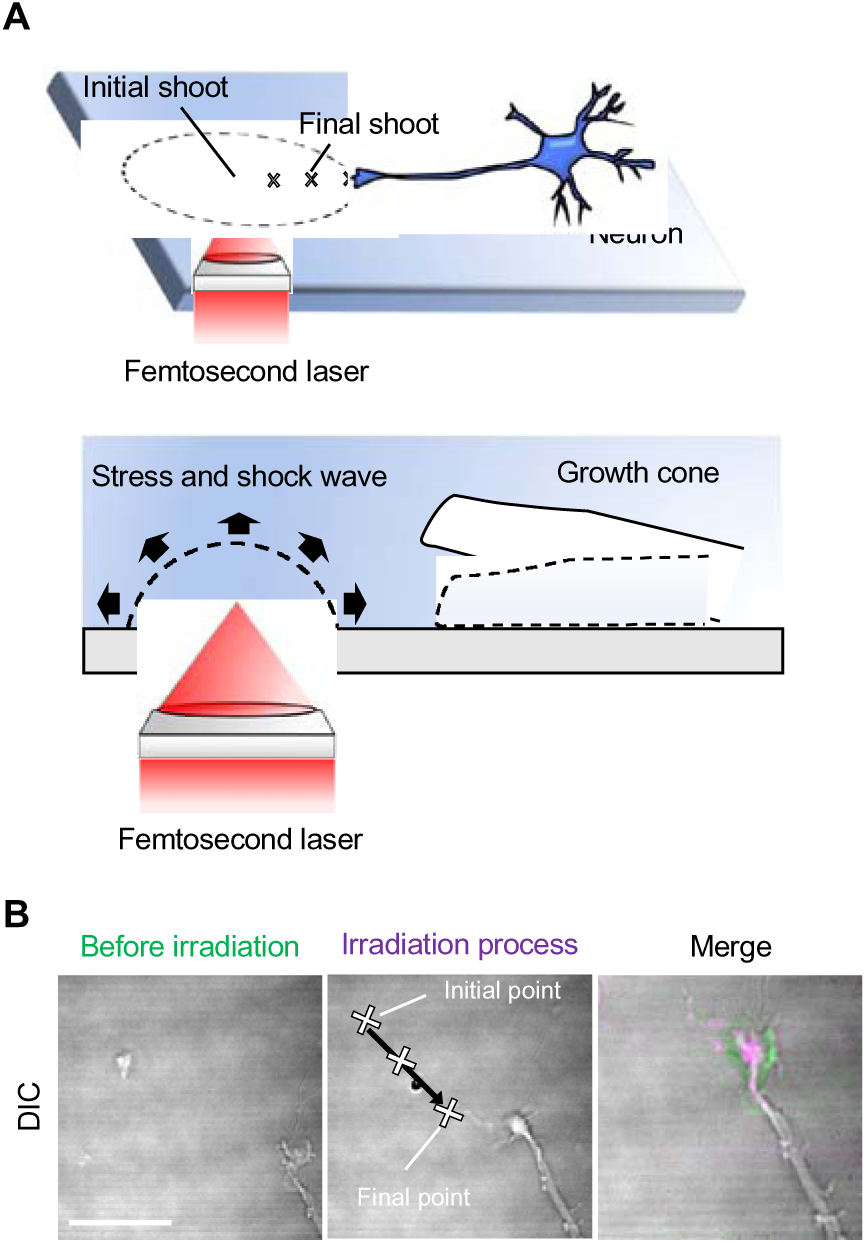
Observation of adhesion breaking of an axonal growth cone by femtosecond laser-induced impulsive force. (A) Schematics of the spatial relation between the femtosecond laser pulse and targeted axonal growth cone of a neuron on a glass substrate. The laser focal point was sequentially moved closer to the growth cone, as indicated by an arrow in the upper cartoon. The effect of the impulsive force to the growth cone was indicated in the lower cartoon, which is enlarged view of the tip of the axon in the upper cartoon. (B) A representative result of adhesion breaking of a growth cone observed by a differential interference contrast (DIC) imaging. The left panel is DIC image before the laser irradiation at the final point. The middle panel indicates the laser irradiation process, in which the laser focal point (cross marks) was sequentially moved along an arrow in the image. The right panel is a merged images of the growth cones before and after the final laser irradiation, which are colorized in green and magenta, respectively. Cross marks are laser focal points. Scale bar, 10 µm.

Representative result of the adhesion breaking of the growth cone is shown in Fig. 1B and Video S1. The growth cone moves on µm scale in the differential interference contrast (DIC) images before and after the laser irradiation at the final laser focal point. Such displacement was only observed at the final laser irradiation point. The distance between the growth cone and the final point became greater with increasing the pulse energy (Fig. 2A). The detachment of the growth cone due to force loading may result from the dissociation of laminin from the substrate. To confirm this possibility, fluorescent dye-conjugated laminins were coated onto the grass-bottom dish, and their fluorescent intensity was monitored following impulsive force loading. As shown in Fig. S1, no significant change in fluorescence intensity was observed around laser focal point. In addition, we have confirmed that the effects of cavitation and temperature are negligible for the cell detachment by the femtosecond laser impulse. These findings suggest that the axonal detachment results from the dissociation of L1 from laminin rather than the detachment of laminin from the substrate. This observation indicates that growth cones can be selectively detached from the substrate by the laser-induced impulsive force.

**Figure 2.**
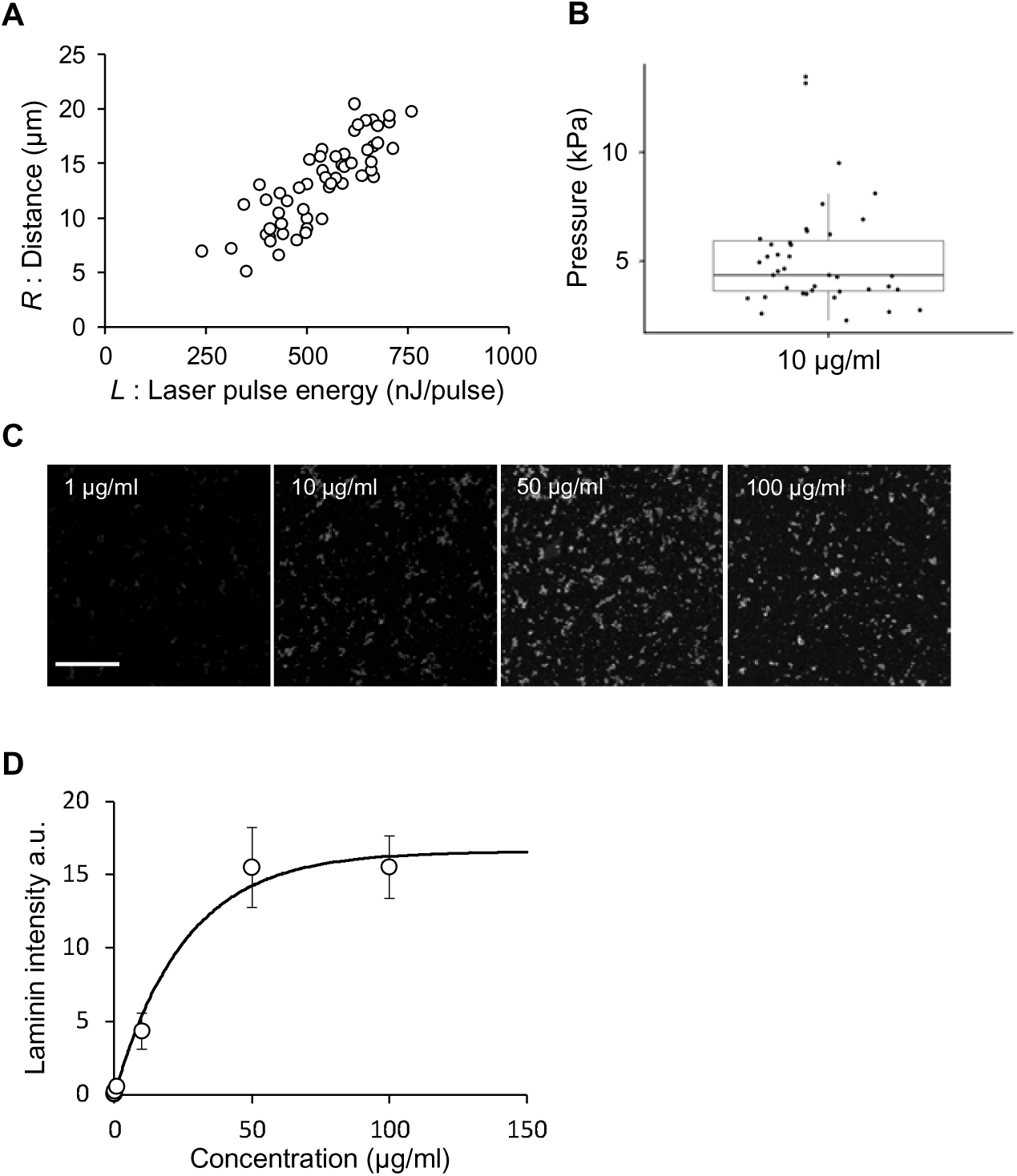
Quantitative evaluation of breaking force for growth cone adhesion by using femtosecond laser impulsive force. (A) Pulse energy *L* dependence of threshold distance *R* to break the growth cone adhesion on a glass substrate coated with a 10 µg/ml laminin solution, corresponding to an *A* of 0.45 (see Eq. [4]). *n* = 36. (B) Box-and-whisker plot of adhesion breaking threshold. Dots in the graph is the threshold calculated independently by substituting data from panel A into Eq. [2]. The mean value and standard deviation are 4.35 and 1.08 kPa, respectively. (C) Images of fluorescent dye-conjugated laminin on the substrate. Concentrations of the laminin solution used for the coating are indicated at the top. Scale bar, 10 µm. (D) Fluorescence intensity as a function of the laminin concentration. The fluorescence intensities were measured on substrates coated with laminin solutions with concentrations of 0.01, 0.1, 1, 10, 50, and 100 µg/ml. The fitting curve was calculated by Eq. [3], where *I*_max_ = 25.5 and *k* = 16.6. *n* = 50 for each concentration. Data are means ± SDs from three independent experiments.

### Quantification of the adhesion breaking force

We evaluated the threshold distance (*R*) to break growth cone adhesion to the glass surface coated with 10 µg/ml laminin at different laser pulse energies (*L*) (Fig. 2A). As the impulsive force near the growth cone increases with *L*, the positive correlation between *R* and *L* indicates that *R* increases with an increasing impulsive force. We quantified the threshold for breaking the adhesion by using our previously established AFM method^24^ in which an AFM cantilever replaces the tip of the growth cone and the impulsive force loaded on the cantilever is estimated from its bending movement^18^ as a function of *L*. From the estimation, the impulsive force *F*_0_ generated at the laser focal point is related to *L* as follows:

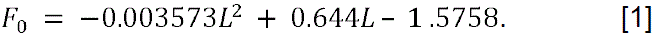

Because the water medium absorbs the pulse with nonlinear processes such as multi-photon and multi-cyclic absorption, the force increases nonlinearly with the pulse energy. Therefore, although the curve in Fig. 2A looks linear in energy, it does not mean that the impulsive force *F*_0_ increases linearly with *R.* Assuming that *F*_0_ propagates spherically in the vicinity of the laser focal point, the impulsive force as a unit of pressure (*P*) is expressed by the following equation:

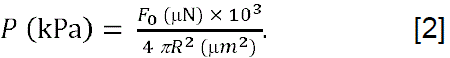

Fig. 2B is a plot of pressure to break the growth cone adhesion calculated with Eq. [2] for each data point in Fig. 2A. The mean value of the minimum pressure to break growth cone adhesion to a 10 µg/ml laminin substrate was 4.35 ± 1.08 kPa, comparable with the breaking threshold reported in our previous study^23,24^.

### Adhesion strength of the growth cone depends on L1CAM-laminin binding

Fluorescent dye-conjugated laminin was used to assess the density of laminin on the glass substrate. The bright spots in images in Fig. 2C are aggregates of laminin less than 1 µm in size. The number of aggregates increased with concentration (*C*) of laminin used to coat the glass and saturated when *C* > 50 µg/ml. The fluorescence was observed not only from the aggregates but also from the substrate. The intensity of fluorescence (*I*) increased with *C* until reaching saturation (Fig. 2D). This result implies that the number of laminin molecules attached to substrate *N* is not proportional to *C* but saturates when *C* > 50 µg/ml. Here we use *I* as an index of coverage of laminin on the substrate. The saturation of *I* is simply expressed by an exponential plateau curve:

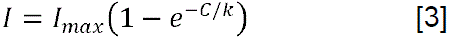

We assumed that (i) *I* is proportional to the number of laminin molecules attached to substrate *N*; (ii) *N* has a maximum that determines *I*_max_. The constant *k* presumably reflects attachment of laminin molecules on the substrate, which are induced by collision of the laminin molecules to the substrate on the coating period (12 h in this experiment). In addition, we neglected the dissociation of laminin from the substrate because *I* was not significantly different after replacing the medium to one without laminin for the laser irradiation experiment. We defined the coverage (*A*) of laminin as an index of laminin density on the substrate using the following equation:

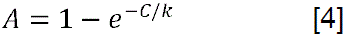

An *A* of 1 means that the laminin attached maximumly on the substrate. The constant *k* was estimated by least-squares fitting with Eq. [3] from the data in Fig. 2D to obtain *A* on the substrate coated with laminin solution at concentration *C*.

The breaking threshold was evaluated according to *A* (left plots in Fig. 3A) and compared with that determined at L1CAM-knockdown samples (right plots in Fig. 3A). The threshold with respect to the coverage of laminin increased with *A* (Fig. 3B). On the other hand, the L1CAM-knockdown samples maintained thresholds that matched those shown by controls at the low coverage (*A* = 0.01). As the expression of L1CAM is almost suppressed by RNAi^12^ (see also Materials and Methods), the major adhesion of the control sample is not due to L1CAM-laminin binding. These findings indicate that the adhesion strength between the growth cone and substrate is strengthened by L1CAM-laminin binding. The offset threshold (∼2.5 kPa) likely reflects laminin-independent adhesive interactions.

**Figure 3.**
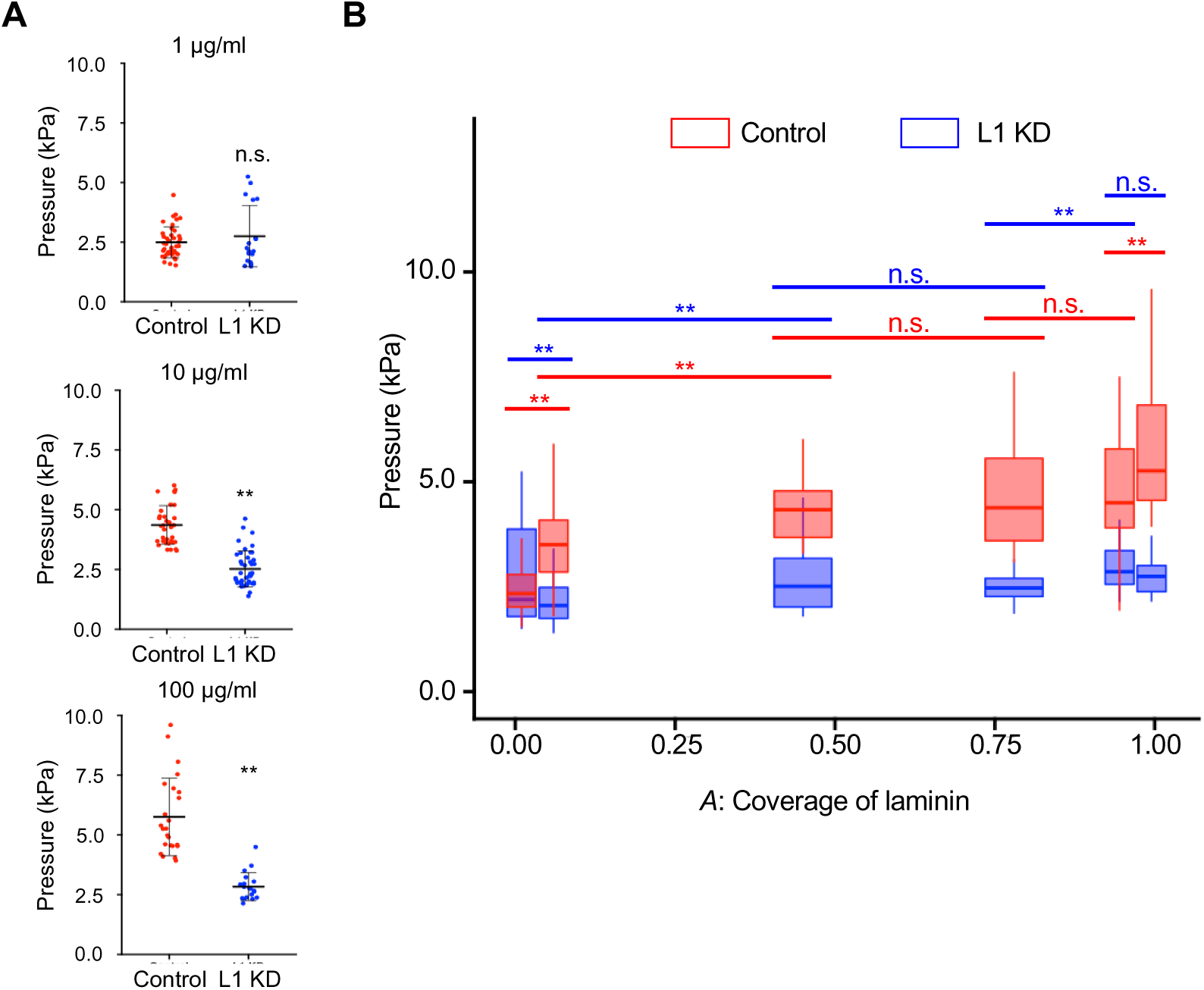
Adhesion breaking of growth cone on a laminin-coated substrate. (A) Means and SDs of the adhesion breaking threshold. Dots in the graph is the threshold calculated independently by substituting data from panel A into Eq. [2]. The laminin concentration is indicated at the top. Red and blue dots indicate control neurons and L1CAM knockdown neurons, respectively. (B) Box-and-whisker plots of adhesion breaking threshold. Control neurons: n = 41, 35, 36, 12, and 25 signals for *A* = 0.01, 0.06, 0.45, 0.78, and 0.99, respectively. L1CAM knockdown neurons: *n* = 18, 44, 34, 20, and 19 signals for *A* = 0.01, 0.06, 0.45, 0.78, and 0.99, respectively. \*\**p* < 0.01 (two-tailed Student’s *t* test), n.s., not significant.

The increase in adhesion strength with increasing *A* in the control samples (Fig. 3B) suggests that the adhesion strength due to the L1CAM-laminin binding increases with the laminin density on the substrate, despite the variability as a result of individual differences among the cells. As the adhesion strength is integral to the individual binding strength between L1CAM and laminin, the adhesion strength reflects the number of L1CAM-laminin interactions. Thus, the increase in the breaking threshold with *A* may reflect the increase in the number of the L1CAM-laminin interactions that occur with increased laminin density. This relationship is presumably satisfied until the L1CAM sites available for binding are saturated.

### L1CAM-laminin binding promotes F-actin-adhesion coupling

The contribution of the laminin coverage *A* to F-actin-adhesion coupling in the retrograde flow was investigated next by visualizing F-actin and L1CAM molecules in filopodia at the growth cone. F-actin dynamics were observed by the motion of fluorescent actin speckles tagged with HaloTag (Fig. 4A; Video S2; Video S3; Video S4) which were observed moving along filaments toward the leading edge of the growth cone. The speed at which they moved decreased linearly with increasing *A*, slowing from 3.70 µm/min at an *A* of 0.06 to 2.29 µm/min at an *A* of 0.99 (Fig. 4B). These data suggest that *A* promotes the cytoskeletal-adhesion coupling.

**Figure 4.**
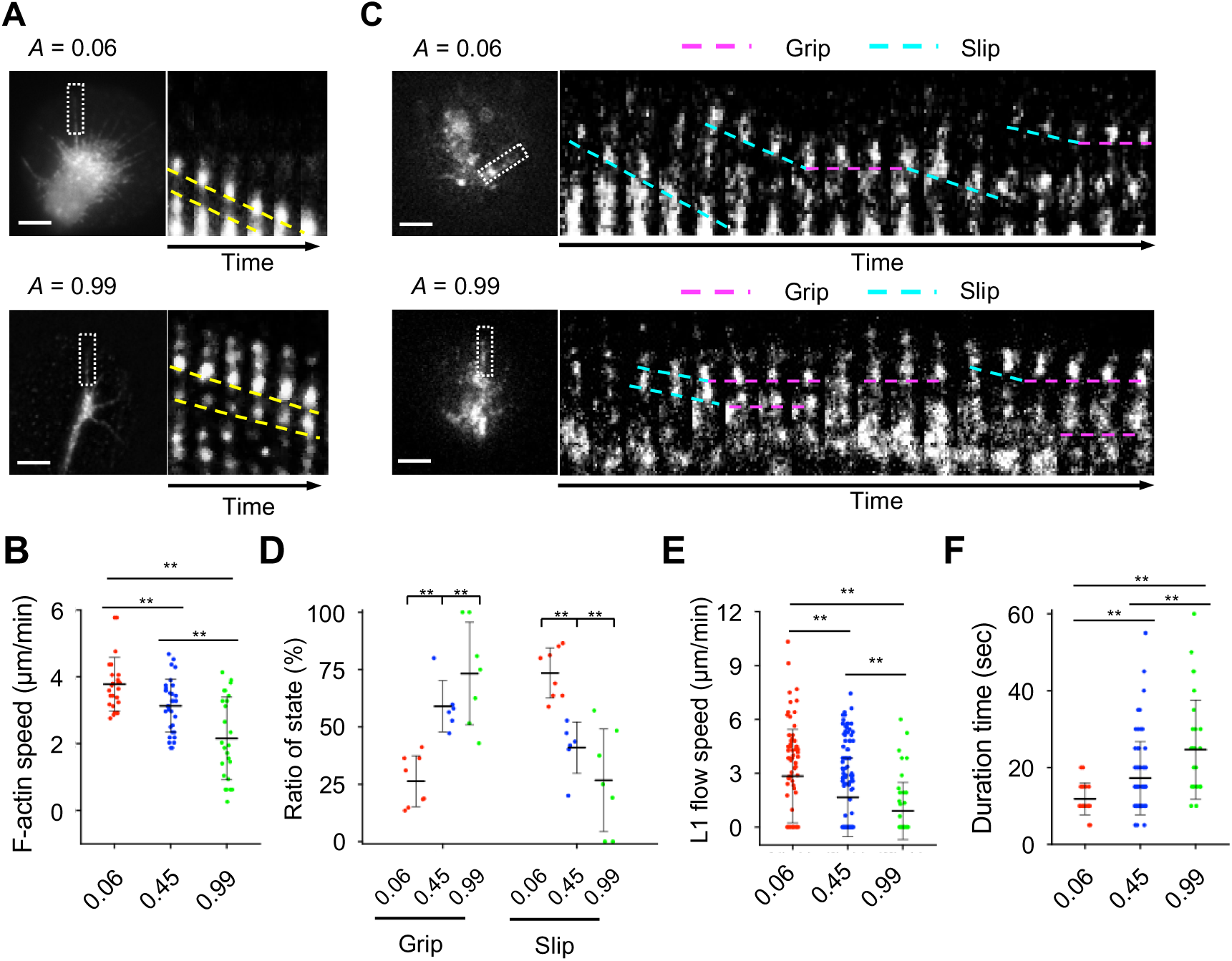
Molecular dynamics of F-actin and L1CAM in the axonal growth cone detected by fluorescence speckle microscopy. (A) Fluorescence speckle images of the HaloTag-actin in a filopodium extended from an axonal growth cone. The coverage of laminin (*A*) is indicated at the top. Kymographs (right) depict HaloTag-actin behavior in boxed region in the image on the left. Slope of the yellow dashed line corresponds to retrograde flow speed of the F-actin. Time interval between frames, 5 s. Scale bar, 5 μm. (B) Retrograde flow speed of F-actin. *n* = 125, 160, and 130 signals for *A* = 0.06, 0.45, and 0.99, respectively. (C) Fluorescence speckle images of L1CAM-HaloTag in a filopodium. Kymographs (right) depict L1CAM-HaloTag behavior in a boxed region in the image on the left. Dashed pink and blue lines connect L1CAM in grip and slip states, respectively. Time interval between frames, 5 s. Scale bar, 5 μm. (D) Ratios of the grip and slip states of L1CAM-HaloTag in filopodia obtained from the kymograph analyses. *n* = 261, 450, and 197 signals for *A* = 0.06, 0.45, and 0.99, respectively. (E) Flow speed of L1CAM-HaloTag in the slip state. The speed corresponds to slopes of dashed blue lines in panel C. (F) Duration time of L1-HaloTag grip. White, red, and blue bars represent data for *A* = 0.06, 0.45, and 0.99, respectively. Data are means ± SDs; \*\**p* < 0.01.

The dynamics of L1CAM-laminin binding were evaluated as grip and slip motions of L1CAM-HaloTag as shown in Fig. 4C (Video S5; Video S; Video S7). The ratios of grip and slip states increased and decreased, respectively, with increasing *A* (Fig. 4D). Consistent with this, the speed at which HaloTag traveled (i.e., flow speed) decreased (Fig. 4E) while the duration spent in the grip phase increased (Fig. 4F) with increasing *A*. The differences between slip and grip states were proportionate to *A* in the range of 0.06 to 0.99.

L1CAM-laminin binding promotes the L1CAM grip state, transmitting the traction force to the substrate^12^. With increased cytoskeletal-adhesion coupling, F-actin flow slows and the traction force transmitted to the substrate for growth cone migration increases^4,25^. Therefore, the associations described above support the result from the adhesion breaking test, i.e., the number of the L1CAM-laminin interactions, reflective of the adhesion strength of the growth cone, are nearly proportional to *A*. Conversely, the dissociation of L1CAM-laminin interactions disrupts cytoskeletal-adhesion coupling such that the force of retrograde flow is no longer transmitted to the substrate^12^. The grip and slip motions observed in this study indicate that L1CAM-laminin binding is not static but changes dynamically. Thus, when the average number of L1CAM-laminin interactions between the growth cone and substrate is increased, the grip state is prolonged.

### L1CAM-laminin binding results in biphasic axonal migration

To examine the effect of laminin coverage on growth cone migration, hippocampal neurons were transfected with either control RNA or L1CAM miRNA, and their migration was monitored over a 7 h in different concentrations of *A* (Fig. 5A left panels; Video S8). The migration speed of the growth cone did not increase linearly but instead exhibited a biphasic response dependent on *A*: within the range of 0.06 to 0.45, the migration speed exhibited a significantly increased, whereas within the range of 0.45 to 0.99, it was markedly reduced (Fig. 5B). In neurons with L1CAM RNAi, the migration speed significantly decreased compared to Control RNAi. Notably, the biphasic migration pattern of the growth cone was abolished in the L1CAM knockdown neurons (Fig. 5A, right panels; Video S8).

**Figure 5.**
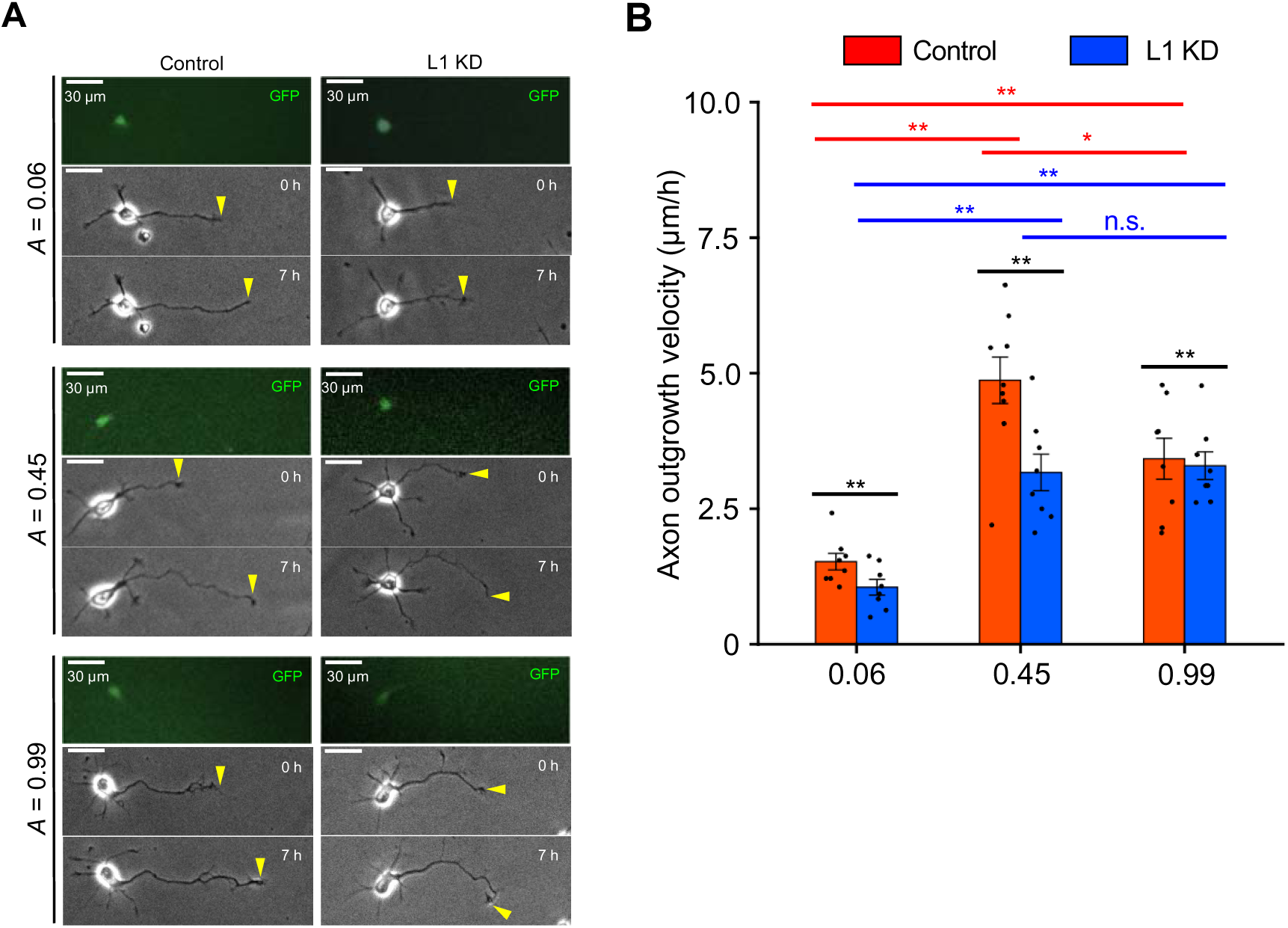
Growth cone migration on laminin-coated substrate. (A) Time-lapse phase-contrast/fluorescence images of hippocampal neurons in control conditions (left panels) or L1 KD conditions (Right panels). Yellow arrowheads indicate growth cone at the first and last time-points, respectively. See also Video – and –. Scale bar: 30 µm. (B) Quantification of axon outgrowth velocity expressing control RNA (red) and L1-CAM miRNA (blue) for *A* = 0.06, 0.45, and 0.99, respectively. Control: *n* = 8, 9, and 8 for *A* = 0.06, 0.45, 0.78, and 0.99, respectively; L1 KD: *n* = 8, 8, and 8for *A* = 0.06, 0.45, and 0.99, respectively. Data are means ± SDs. Kruskal-Walls test. \**p* < 0.05, \*\**p* < 0.01. n.s., not significant.

### L1CAM-laminin binding results in biphasic axon outgrowth

Axon outgrowth also depends on the laminin coverage (*A*), as shown in Fig. 6. Axon lengths were the longest (>130 µm) when neurons were cultured under conditions where *A* was between 0.78 and 0.95. However, the lengths of axons from L1CAM knockdown neurons were not affected by *A* (blue box in Fig. 5B), with shorter axons overall. These data suggest that axon outgrowth is regulated by the L1CAM-laminin binding to the laminin substrate and that the regulation is biphasic.

**Figure 6.**
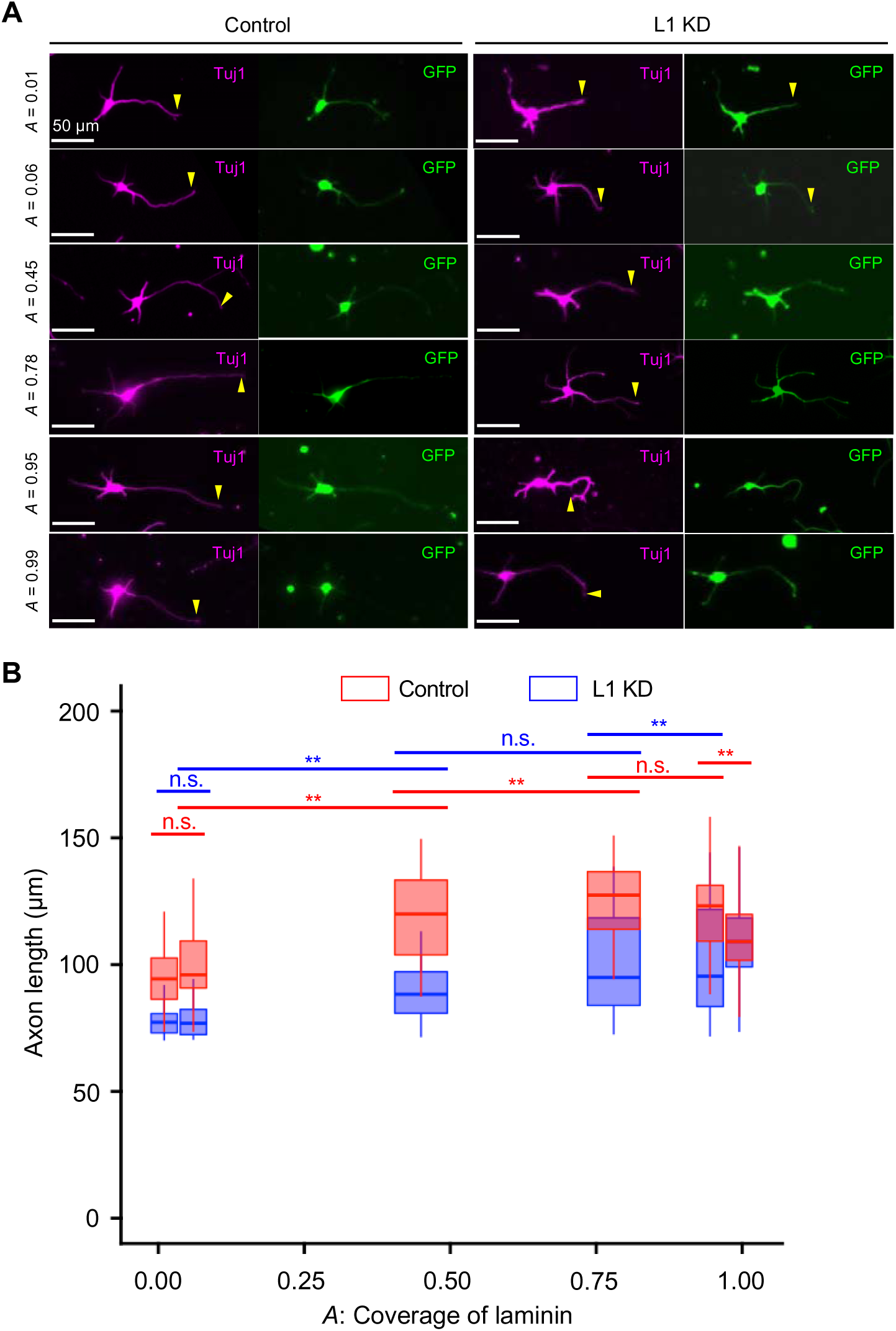
Axon elongation on laminin-coated substrate. (A) Confocal images of neurons visualized with a GFP (green) and Tuj1 (Tubulin β3, magenta) antibody. Neurons were cultured on polyacrylamide gels coated with laminin. The coverage of laminin (*A*) is indicated on the left. Scale bar; 50 µm. (B) Box-and-whisker plots of axon length. Control neurons (red): *n* = 91, 28, 79, 60, and 71 signals for *A* = 0.01, 0.06, 0.45, 0.78, and 0.99, respectively; L1CAM knockdown neurons (blue): *n* = 43, 49, 73, 103, and 48 signals for *A* = 0.01, 0.06, 0.45, 0.78, and 0.99, respectively. Kruskal-Walls test. \**p* < 0.05, \*\**p* < 0.01. n.s., not significant.

### Traction force for the axon outgrowth exhibits biphasic behavior

Traction force microscopy was utilized for monitoring the force loaded on the substrate in the axon outgrowth. In this method, the traction force is observed as displacement of the fluoresce microbeads enclosed in the substrate and quantified from the bead shift from original position to displaced position in the axon outgrowth (Fig. 7A). At the region of the growth cone, the mean force for *A* = 0.45 (35 pN/nm^2^) was larger than not only that for *A* = 0.01 (25 pN/nm^2^) but also that for *A* = 0.99 (25 pN/nm^2^) (Fig. 7B). The dependence, which is not monophasic but biphasic, is similar to that for axon outgrowth (Fig. 5), suggesting that the traction force is directly related with the axon outgrowth.

**Figure 7.**
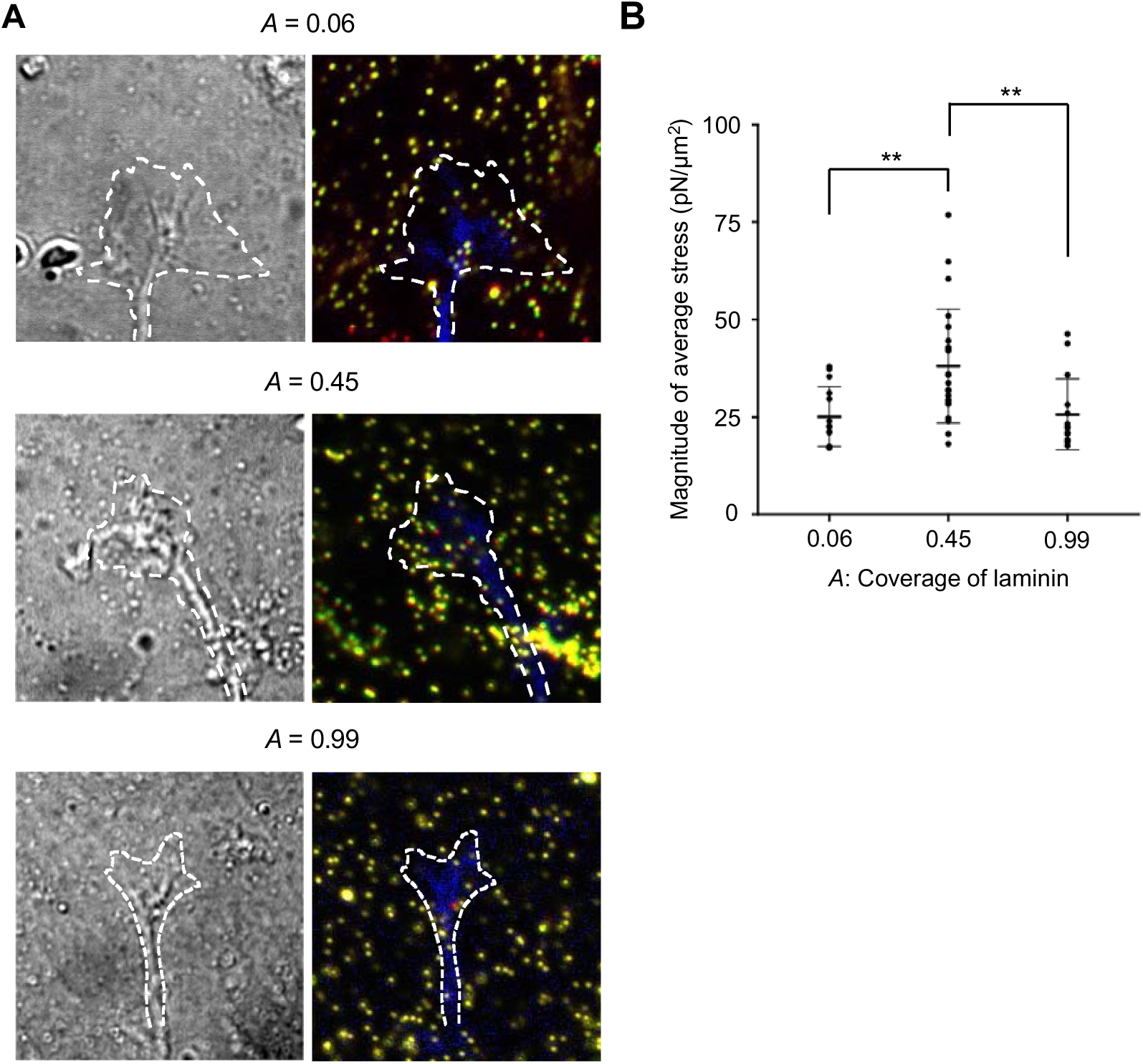
Measurement of traction force on a laminin-coated substrate. (A) DIC (left panels) and fluorescence images (right panels) of an axonal growth cone cultured on polyacrylamide gels with embedded 200 nm fluorescent beads. Dashed lines indicate the boundary of the growth cone. The blue color in the dashed line is due to EGFP which is expressed in the neuron to track the growth cone position. The original and displaced bead positions in the gel are indicated by green and red colors in fluorescence images, respectively. The yellow color overlaps the green and red colors, i.e. the bead is rarely displaced. Laminin coverage *A* is shown at the top of the panel column. Scale bar; 10 µm. (B) Magnitude of the traction forces under axonal growth cones. Data represent mean ± SEM (error bars). Kruskal-Walls test. \*\**p* < 0.01.

## Discussion

In this study, we quantified adhesion strength of axonal growth cone (Fig. 3B), axon elongation (Fig. 5, 6), and traction force of growth cone in the elongation (Fig. 7B) as a function of laminin density on the substrate. In addition, we evaluated molecular dynamics of F-actin and L1CAM in the growth cone (Fig. 4). Although the adhesion strength increased with the laminin density, the laminin dependency of the axon elongation and traction force indicated biphasic behavior. These differences revealed in the measurements allow us to argue the mechanism of axonal outgrowth promoted by the growth cone.

The adhesion strength measured by femtosecond laser impulses is not only for the adhesion between L1CAM and laminin but for total amount of the adhesion including other specific and nonspecific adhesions. The adhesion strength estimated as the adhesion breaking force is on the order of kilopascals (3-5 kPa). On the other hand, traction force detects the force to promote axon outgrowth. The force is generated by force transmission of F-actin retrograde flow; transmission is effective when the binding forces of the clutch and adhesion molecules between F-actin and substrate are higher than the traction force^12^. The force is estimated in the range 20 – 50 pN/μm^2^. Since the unit of force for traction (pN/μm^2^) corresponds to pascals (Pa = N/m^2^), the adhesion strength is about 100 times larger than the traction force (0.02 – 0.05 kPa). As shown in Fig. S1, the laser impulse does not directly break the laminin on the substrate. This means the adhesion breaking force in Fig. 3 reflects the adhesion between the growth cone and the substrate. The traction force in Fig. 7 is the force transmitted by the F-actin retrograde flow. Conclusively, our results suggest that the adhesion between the growth cone and the substrate is strong enough to transmit the force of F-actin retrograde flow to the substrate.

The L1CAM-laminin interactions promote the axon outgrowth transmitting the force of F-actin retrograde flow to the substrate^26^. In this study, we confirmed that axon outgrowth depends on laminin concentration, incorporating adhesion strength and traction force. When the laminin density is low, F-actin is not coupled to the adhesive substrate through clutch molecules (e.g., shootin1a and cortactin) and L1CAM. As a result, the force of retrograde flow is not effectively transmitted to the substrate to produce sufficient traction for axon outgrowth. This was demonstrated by the short axon lengths of neurons cultured on the substrate with low laminin density. By contrast, the rate of retrograde flow slows as the laminin density increases, indicating greater cytoskeletal-adhesion coupling that promotes transmission of the force from the flow to the traction force for growth cone migration. Our results indicate that laminin densities resulting in an *A* between 0.45 and 0.78 are optimal for providing suitable traction force. Interestingly, we observed a decrease in axon outgrowth at a high laminin density, suggesting that excessive L1CAM-laminin binding suppresses axon outgrowth.

Similar biphasic behavior has been reported depending for concentration of guidance molecules^17,27^ and integrin-ligand binding^28–30^. Notably, neurite outgrowth exhibits a biphasic dependence on ligand concentration, with intermediate concentrations resulting in maximal neurite outgrowth^31^. Our results consistent with previous studies, reinforcing the validity of our method. By integrating genetic manipulation, we extend mechanical measurement to provide the specific contribution of L1-laminin interactions to adhesion strength. Moreover, our study extends these findings by demonstrating that traction force generation also exhibits a biphasic dependence with laminin concentration. This integration of force measurement and genetical approaches helps fill the gap between classical adhesion studies and contemporary molecular insights into growth cone migration.

Cell adhesion strength was also evaluated by shear-stress flow assay and compared with the migration speed. With strong adhesion, the cells spread and extended lamellae, but the cell body did not move. The suppression of migration was attributed to the inability of cells to overcome adhesion to the substrate^30,32–34^. For axons, outgrowth is promoted not only by the traction force at the forward side but also by detachment of the back side. Thus, the decrease in growth we observed may be attributable to a lack of detachment as a result of excessive L1CAM-laminin interactions which does not couple with the retrograde flow. On this assumption, the traction force is also suppressed by the excessive interaction as well as the axon outgrowth.

Notably, axon outgrowth was suppressed when *A* increased from 0.78 to 0.99, an estimated proportional increase in adhesion strength of ∼20%. This result indicates that the modification of outgrowth is on the order of 10% of the modification in the adhesion strength as a result of the number of L1CAM-laminin interactions. Our data therefore suggest that these interactions are responsible for adhesion to the substrate and thus for the regulation of axon outgrowth This regulation is key for growth cone migration and axon outgrowth through the extracellular matrix in brain, thereby contributing to the formation of network connections with other neurons and relevant cells.

This investigation utilized femtosecond laser impulses to quantitatively evaluate the adhesion strength between axonal growth cones and a laminin substrate. The data show relative contribution of the L1CAM-laminin interactions in adhesion of the growth cone i.e., the strength of the interaction increases with the laminin density. Notably, axon outgrowth does not increase monotonically with increased L1CAM-laminin binding but instead exhibits biphasic behavior, in which outgrowth is suppressed in the presence of high amounts of L1CAM-laminin binding. This biphasic outgrowth is regulated by altered adhesion caused by changes in the number of binding interactions on the order of 10%. These results suggest that the balance between the traction force from the cytoskeletal-adhesion coupling and growth cone adhesion is one of the keys to regulating axon outgrowth. Future studies on the biphasic regulation of axon outgrowth should seek to further elucidate the guidance mechanism.

## Materials and Methods

### Preparation of cell culture substrate

For each experiment, a 35 mm glass-bottom dish (Matsunami, Osaka, Japan) was coated with 100 µg/ml poly-lysine (FUJIFILM WAKO Pure Chemical Corporation) at 37°C for 12 h. After washing, the plate was coated with laminin (laminin 1; FUJIFILM WAKO Pure Chemical Corporation) in phosphate-buffered saline (PBS) at 37°C for 12 h. The surfaces were washed three times with PBS. The laminin density on the dish was modified by altering the concentration of the deposited laminin solution (0.01–100 µg/ml). The laminin density was evaluated by coating the dish with laminin conjugated to a fluorescent dye (green fluorescent HiLyte 488; Cytoskeleton), which was observed under a confocal laser scanning microscope (Zeiss LSM710; excitation, 488 nm; emission, 510 nm). The fluorescence intensity was estimated as an area integration (15 × 15 µm) of the substrate fluorescence.

### Cell culture

Hippocampal neurons were prepared from embryonic day 18 rats and seeded on glass-bottom dishes. To induce axon outgrowth, neurons were cultured in neurobasal medium (Thermo Fisher Scientific) containing B-27 supplement (Thermo Fisher Scientific) and 1 mM glutamine for 3 days. All relevant aspects of the experimental procedures were approved by the Institutional Animal Care and Use Committee of Nara Institute of Science and Technology.

### Femtosecond laser irradiation system

The cultures were imaged on an inverted microscope (IX71; Olympus) utilizing femtosecond laser pulses from a regeneratively amplified Ti:Sapphire femtosecond laser (800 ± 5 nm, 100 fs, <1 mJ/pulse, 32 Hz) (Solstice Ace; Spectra-Physics). The pulse was focused near the growth cone (Fig. 1A) through a 100× lens objective (UMPlanFl, numerical aperture [NA], 1.25; Olympus). The irradiation was controlled by a mechanical optical shutter (Σ-65GR; Sigma Koki). The laser pulse energy was tuned by a half-wave (λ/2) plate and dual polarizers. A single femtosecond pulse (50–1,000 nJ/pulse) was applied near the growth cone, and adhesion breaking was monitored by a charge-coupled-device (CCD) camera.

### Impulsive force measurement system using AFM

AFM was used to quantify the force needed to break the adhesion, as described previously^18^. An AFM cantilever (thickness, 4.0 µm; length, 125 µm; width, 30 µm; force constant, 42 N/m; resonance frequency, 330 Hz in air) (TL-NCH-10; Nano World, Neuchatel, Switzerland) was attached to the AFM head (Nano Wizard 4 BioScience; JPK Instruments, Berlin, Germany) and placed in pure water on the microscope stage. The laser pulse was focused 10 µm away from the top of the cantilever. The transient oscillation of the cantilever induced by the laser pulse irradiation was detected by an oscilloscope. The magnitude of the cantilever movement was estimated from the oscillation.

### Evaluation of laminin detachment on the substrate by impulsive force loading

The system for the laser irradiation and the condition of fluorescence laminin-coat are the same as described above. The change in fluorescence intensity before and after laser irradiation was analyzed in estimated shock wave generation area.

### L1CAM knockdown experiment

L1CAM knockdown neurons were prepared by using a Block-iT Pol II miR RNAi expression kit (Invitrogen). The targeting sequence of L1CAM miRNA and its effectiveness were reported previously^12^. Hippocampal neurons were transfected with the miRNA expression vector and incubated for 20 h. The cells were then collected and cultured on the laminin-coated glass-bottom dishes. In this system, GFP is expressed with the L1CAM miRNA, enabling the growth cones of transfected cells to be visualized and monitored.

### Fluorescent speckle microscopy

The retrograde flow of F-actin and slip and grip motions of L1CAM were investigated by fluorescent speckle microscopy. HaloTag-actin and L1CAM-HaloTag were expressed in hippocampal neurons. To introduce HaloTag tetramethylrhodamine (TMR) to L1CAM-HaloTag and HaloTag-actin, hippocampal neurons were incubated with HaloTag TMR ligand (Promega) at a 1:1,500 dilution in L15 medium containing B27 supplement and 1 mM glutamine for 1 h at 37°C. The medium was then replaced with fresh L15 medium. The preparation method of HaloTag-actin is specified in the literature^12^.

HaloTag-actin speckles were observed at 37°C using a fluorescence inverted microscope (Axio Observer A1; Carl Zeiss) equipped with a C-apochromat 63× NA 1.20 lens objective (Carl Zeiss), an illumination laser (561 nm), and an EM-CCD camera (Ixon3; Andor). Fluorescent L1CAM-HaloTag speckles in growth cones were observed using total internal reflection fluorescence (TIRF) on an inverted microscope (IX81; Olympus) equipped with a TIRF lens objective (UAPON 100×OTIRF NA 1.49; Olympus), an illumination laser (488 nm), and a scientific complementary metal-oxide semiconductor (sCMOS) camera (ORCA Flash4.0LT; HAMAMATSU). The flow speed of F-actin and slip speed of L1CAM were analyzed by monitoring the fluorescence signals of the HaloTags at 5 s intervals. L1CAM puncta that were visible for at least 10 s (two intervals) were analyzed; immobile ones were defined as L1CAM in stop (grip) phase, while those that flowed retrogradely were defined as in flow (slip) phase.

### Evaluation of growth cone migration by confocal microscopy

In the neuron, EGFP is expressed to track the growth. The growth cone was monitored by the EGEP fluorescence and DIC images. The surface of polyacrylamide gels was coated with laminin using the same procedure as that used to coat glass-bottom dishes (see above). EGFP and DIC signals were observed at 5 min intervals for 7 h using a fluorescence microscope (IX81; Olympus) equipped with an EM-CCD camera (Ixon DU888; Andor), using a 20×/0.8 objective lens. Speed of growth cone migration was measured using ImageJ (Fiji version).

### Evaluation of axon length by immunofluorescence staining

Axon length was evaluated by immunofluorescence imaging. Neurons were cultured on polyacrylamide gels coated with laminin for 3 days, fixed with 3.7% formaldehyde in PBS overnight at 4°C, treated for 15 min with 0.05% Triton X-100 in PBS at 4°C, and then incubated with 10% fetal bovine serum in PBS overnight at 4°C. The cells were then incubated with anti-GFP antibody (MBL, 598) and anti-Tubulin β3 (Biolegend, 801201), as described by Toriyama et al^25^. We used secondary antibodies conjugated with Alexa Fluor 488 (Invitrogen, A11008) or Alexa Fluor 594 (Invitrogen A11032), and observed with a confocal laser scanning microscope (LSM 710). The lengths of all axons of 50 cells were measured for each coverage.

### Traction force microscopy

Traction force microscopy was performed as described in literatures^12,25^. Briefly, neurons were cultured on polyacrylamide gels with embedded fluorescent microspheres (200 nm diameter; Invitrogen). Time-lapse imaging of florescent microsphere and growth cones was performed at 37℃ using the confocal microscope (LSM710; Carl Zeiss) with a C-Apochromat 63x/1.2 W Corr objective (Carl Zeiss). In the neuron, EGFP is expressed to track the growth. The growth cone was monitored by the EGEP fluorescence and DIC images. The surface of polyacrylamide gels was coated with laminin using the same procedure as that used to coat glass-bottom dishes (see above).

Traction forces under the growth cones were detected by displacement of the beads from their original positions. Force vectors were estimated from the displacement. To compare the forces under different laminin densities, the magnitude of the force vectors of individual beads under the growth cones were statistically analyzed and expressed separately as means ± SEM. The quantification of traction force was performed using MATLAB code TFM2021 and software^35^.

### Statistical analysis

Graphs were generated and statistical analyses were performed by using Microsoft Excel and R. For nonparametric datasets, we used Mann–Whitney test to compare two groups, and Kruskal–Wallis test with Dunn’s multiple comparison to compare more than two groups.

## Acknowledgments

We thank Dr. Ryohei Yasukuni and Dr. Kazunori Okano (Nara Institute of Science and Technology, Japan) for fruitful discussions.

## Author Contributions

S.Y. and Y.H. designed the research. N.I. supervised the research and contributed to presentation of the mechanism of axon outgrowth. S.Y. performed almost all experiments and data analysis. K.B. prepared primary cultured neurons. S.Y. and Y.H. wrote the article.

