## supplementary Figure for "Cell Adhesion-Dependent Biphasic Axon Outgrowth Elucidated by Femtosecond Laser Impulse"

### Supplementary Information for
Cell Adhesion-Dependent Biphasic Axon Outgrowth Elucidated by Femtosecond Laser Impulse

Sohei Yamada[a,b,†,*], Kentarou Baba[c,†], Naoyuki Inagaki[c], Yoichiroh Hosokawa[a,d,*]

Paste the full author list here

**This PDF file includes:**

Legends for Video S1 to 8

**Other supplementary materials for this manuscript include the following:**

Video S1 to 8

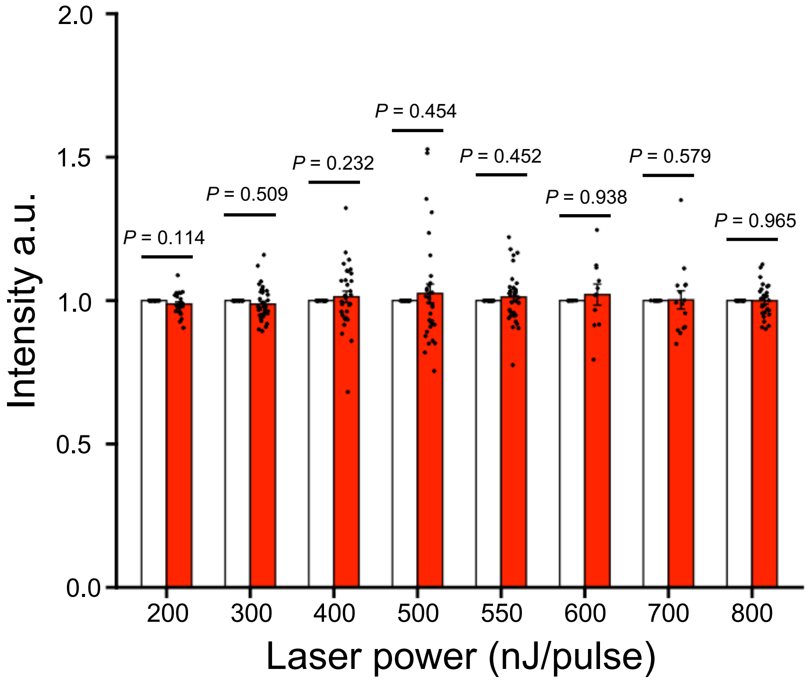


**Supplementary Figure 1. Effect of femtosecond laser-induced impulse loading on laminin.** Quantification of HiLyte 488-conjugated laminin intensity before and after impulsive force loading. 200 nJ/pulse: n = 26, 300 nJ/pulse: n = 32, 400 nJ/pulse: n = 34, 500 nJ/pulse: n = 32, 550 nJ/pulse: n = 32, 600 nJ/pulse: n = 15, 700 nJ/pulse: n = 11, 800 nJ/pulse: n = 26. One-way ANOVA followed by post hoc Tukey-Kramer test. *P* values are indicated in figure. Data are presented as mean ± SD.

**Supplemental Video Legends**

**Video S1.** Adhesion breaking of axonal growth cone by femtosecond laser-induced impulsive force. Time-lapse movie of axonal growth cone of hippocampal neuron cultured on 10 µg/ml laminin (see Fig. 2D). White dot indicates laser focal point. Adhesion breaking of an axonal growth cone was monitored at 1 s intervals.

**Video S2.** Movement of fluorescent features of HaloTag-actin in an axonal growth cone of a neuron cultured on laminin at an *A* of 0.06 (see Fig. 4A top). Images of HaloTag-actin in the growth cone were captured using a fluorescence microscope (Axioplan2; Carl Zeiss). The axonal growth cone was monitored at 5 s intervals.

**Video S3.** Movement of fluorescent features of HaloTag-actin in an axonal growth cone of a neuron cultured on laminin at an *A* of 0.45. Images of HaloTag-actin in the growth cone were captured using a fluorescence microscope (Axioplan2; Carl Zeiss). The axonal growth cone was monitored at 5 s intervals.

**Video S4.** Movement of fluorescent features of HaloTag-actin in an axonal growth cone of a neuron cultured on laminin at an *A* of 0.99 (see Fig. 4A bottom). Images of HaloTag-actin in the growth cone were captured using a fluorescence microscope (Axioplan2; Carl Zeiss). The axonal growth cone was monitored at 5 s intervals.

**Video S5.** Movement of fluorescent features of L1CAM-HaloTag in an axonal growth cone of a neuron cultured on laminin at an *A* 0.06 (see Fig. 4C top). Images of L1CAM-HaloTag in the growth cone were captured using a TIRF microscope (IX81; Olympus). The axonal growth cone was monitored at 5 s intervals.

**Video S6.** Movement of fluorescent features of L1CAM-HaloTag in an axonal growth cone of a neuron cultured on laminin at an *A* of 0.45. Images of L1CAM-HaloTag in the growth cone were captured using a TIRF microscope (IX81; Olympus). The axonal growth cone was monitored at 5 s intervals.

**Video S7.** Movement of fluorescent features of L1CAM-HaloTag in an axonal growth cone of a neuron cultured on laminin at an *A* of 0.99 (see Fig. 4C bottom). Images of L1CAM-HaloTag in the growth cone were captured using a TIRF microscope (IX81; Olympus). The axonal growth cone was monitored at 5 s intervals.

**Video S8.** Migrations of hippocampal neuron expressing control miRNA or L1CAM miRNA cultured on different *A* of Laminin. DIC images of the growth cones were captured using a fluorescence microscope (IX81; Olympus). The axonal growth cone was monitored at 5 min intervals.
